# MRS4Brain: a processing toolbox for preclinical MR spectroscopy and spectroscopic imaging data

**DOI:** 10.64898/2025.12.12.693881

**Authors:** Brayan Alves, Tan Toi Phan, Guillaume Briand, Alessio Siviglia, Gianna Nossa, Jessie Mosso, Eloïse Mougel, Jamie Near, Thi Ngoc Anh Dinh, Omar Zenteno, Bernard Lanz, Thanh Phong Lê, Cristina Cudalbu

## Abstract

**Objectives:** Magnetic resonance spectroscopy is a non-invasive technique for probing metabolism and underpins advanced methods such as magnetic resonance spectroscopic imaging (MRSI) and diffusion-weighted spectroscopy (DWS). MRSI enables spatial mapping of metabolite distributions, offering insights into regional metabolic heterogeneity that single-voxel spectroscopy (SVS) cannot capture. However, MRSI produces large multidimensional datasets and requires complex processing pipelines, limiting reproducibility and accessibility. While human studies benefit from advanced processing tools, similar developments in preclinical research remain scarce, highlighting a demand for practical tools accessible to non-experts.

**Methods:** To address this need, we introduce the *MRS4Brain Toolbox*, a freely available MATLAB-based platform for preclinical spectroscopy, including MRSI, SVS, and DWS.

**Results:** The toolbox integrates reconstruction, preprocessing, quantification, quality control, brain segmentation automatically overlaid on metabolite maps, modeling, and statistical analysis into unified workflows accessible via a graphical interface.

**Conclusion:** By streamlining data processing and reducing technical barriers, *MRS4Brain Toolbox* promotes reproducibility, harmonization, and broader adoption of advanced spectroscopic techniques in preclinical studies, ultimately facilitating translational research.

## 1. Introduction

Magnetic resonance spectroscopy (MRS) is a non-invasive, versatile, and well-established technique for studying metabolism. It serves as the foundation for magnetic resonance spectroscopic imaging (MRSI) and advanced approaches such as diffusion-weighted spectroscopy (DWS). MRSI builds upon single-voxel spectroscopy (SVS) by enabling spatial mapping of metabolite distributions within a single acquisition, thereby providing insights into regional metabolic heterogeneity. This spatial dimension is particularly important as metabolic alterations often exhibit localized patterns that SVS alone cannot adequately capture. With the ability to simultaneously acquire spectra in different locations, MRSI has become a key player in numerous human studies, such as research on glioma or sclerosis [1–5], and has recently attracted interest in preclinical research [6, 7]. These advancements underscore the increasing recognition of MRSI as a powerful modality for understanding the underlying metabolism in disease progression.

Despite its advantages, MRSI acquisitions inherently produce large, multidimensional datasets, as the number of spectra scales with the phase-encoding steps. This complexity necessitates robust visualization and interpretation strategies to ensure reproducibility and clarity. A major challenge common to all MRS methods is the complexity of the data processing pipeline, which requires extensive knowledge of acquisition and multiple preprocessing steps. These steps often involve different tools to achieve a thorough and effective analysis [8, 9]. As methodological advances continue to push the boundaries of MRS capabilities, the complexity of data processing increases, introducing heterogeneities across studies and making harmonization even more difficult. Without unified workflows, these challenges can hinder reproducibility and slow down the adoption of advanced techniques.

Recent consensus on MRS has provided a framework for standardizing essential steps for spectroscopic data analysis quantification [10–12]. Consequently, substantial efforts have been made in the human research realm, where the emergence of MRS tools, such as Osprey [9], FID-A [10], FSL-MRS [8], Gannet [13], ORYX-MRSI [14], and MRSpa [15], has led to the wider spread of the MRS methodologies. To the best of our knowledge, such advanced developments remain scarce in the preclinical research domain. With the recent advent of free induction decay (FID)-MRSI in preclinical research [6], there is demand for practical tools that enable harmonized analyses and facilitate dissemination to non-MRSI experts. Bridging this gap is critical for accelerating translational research and aligning preclinical findings with clinical studies.

This paper introduces the *MRS4Brain Toolbox*, a freely available MATLAB-based platform specifically designed for Bruker preclinical spectroscopy studies. It offers automatic workflows via a graphical user interface for MRSI, SVS, and DWS, aiming to enhance accessibility for researchers starting the preclinical spectroscopy field. By integrating reconstruction, preprocessing, quality control, quantification, modeling, brain segmentation automatically overlaid on metabolite maps, and statistical analysis into a unified environment, the toolbox streamlines complex processing steps and supports comprehensive analysis of preclinical datasets. Ultimately, *MRS4Brain Toolbox* seeks to promote reproducibility, reduce technical barriers, and foster broader adoption of advanced spectroscopic techniques in preclinical research.

## 2. Methods

The current version of the *MRS4Brain Toolbox* is available on the GitHub repository [16], providing functionalities for MRSI, SVS, and DWS (**Figure 1**). Detailed methods are described in the following sections, while the installation guidelines are provided in the Supplementary Information.

**Figure 1.**
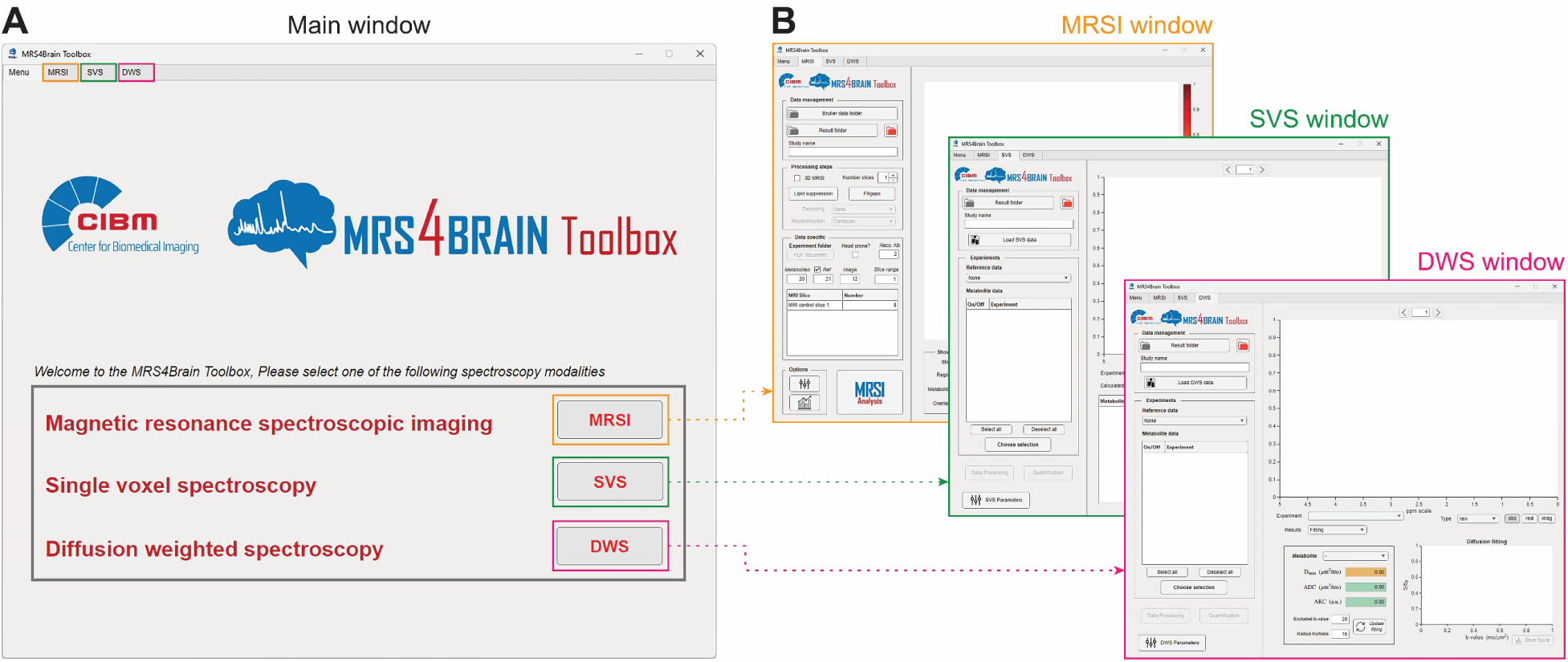
Overview of the MRS4Brain Toolbox. (**A**) Main window of the toolbox (**B**) Graphic user interfaces for MRSI (window highlighted in orange), SVS (window highlighted in green), and DWS (window highlighted in pink).

### 2.1. MRSI processing pipeline

#### 2.1.1. Processing methods

The MRSI processing pipeline (**Figure 2A**) takes magnetic resonance imaging (MRI) images (here, T_2_-weighted coronal images), metabolite, and water MRSI data as inputs. The toolbox can effectively read data reconstructed and provided by various versions of Bruker’s ParaVision 360 software. MRI data needs to be in *NIFTI* format (*\*.nii* or *\*.nii.gz*), while MRSI data can be read as a *\*.fid* file (versions v1.x to v2.0) or a *\*.fid_proc.64* (versions v3.0 and above). Data selection can be managed in the Data Management and the *Data-specific* section of the MRSI window.

**Figure 2.**
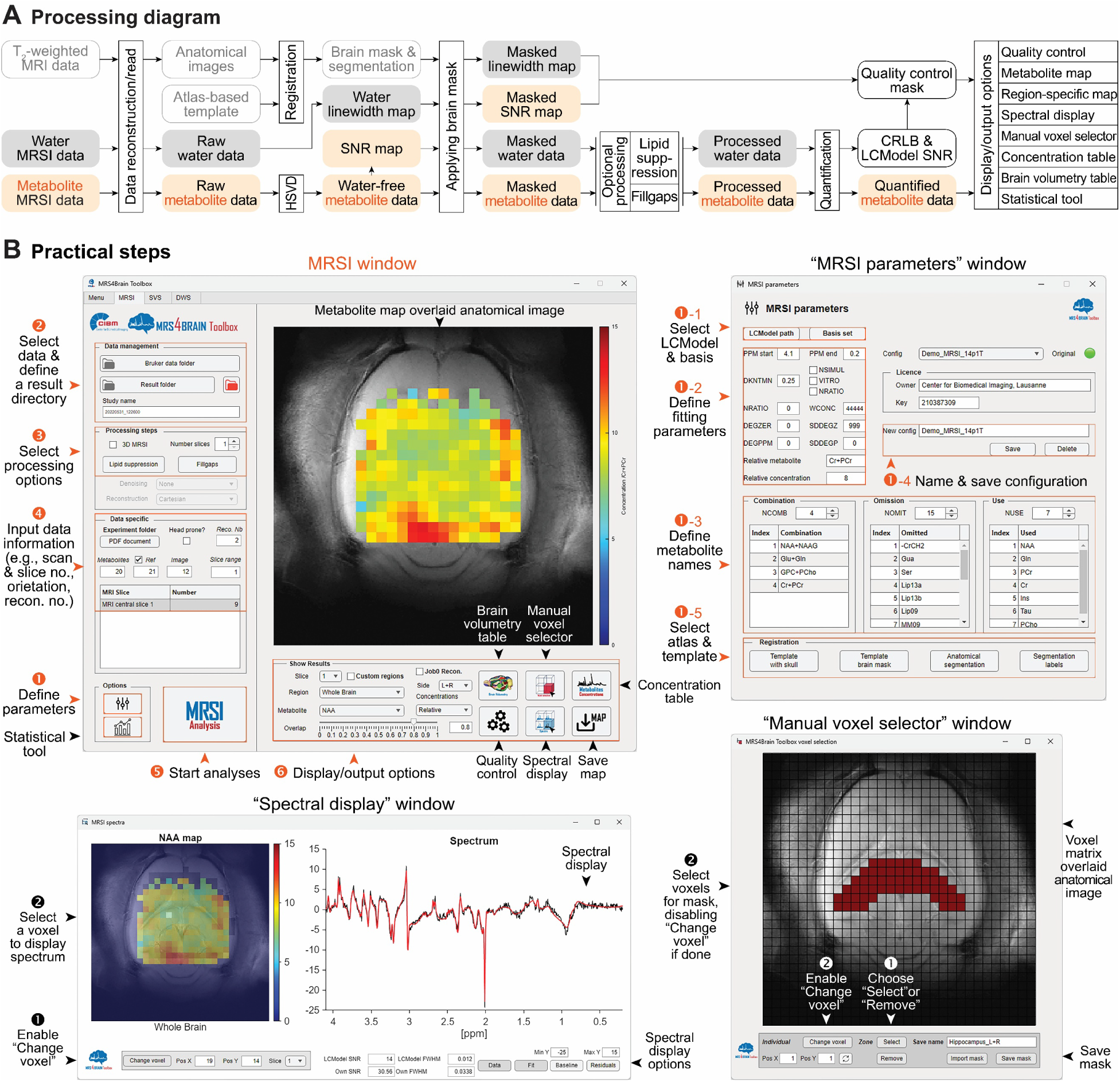
Processing pipeline and practical steps for MRSI data. (**A**) Processing pipeline diagram from data reconstruction, processing to result display, and outputs. (**B**) Typical workflow for processing an MRSI dataset. Top-left: Main MRSI window displays steps 1–6, guiding users through parameter configuration, data input, processing option selection, and display/output choices. Top-right: *MRSI Parameters* window (step 1) for defining parameter configurations. Bottom-left: *Spectral display* window for visualizing spectra in individual voxels. Bottom-right: *Manual voxel selector* for defining user-specified regions of interest.

##### Co-registration

One key motivation for combining anatomical information from MRI data is the ability to segment the different brain regions and generate precise anatomical masks. These masks can then be applied to the MRSI dataset to examine the metabolite contents within specific brain regions. To achieve this, a co-registration between a template with atlas-based segmentation and acquired MRI images is done using Advanced Normalization Tools (ANTs) [17]. This process employs rigid, affine, and symmetric normalization transformation methods in ANTs to align the template to the MRI image, yielding a transformation matrix that is used to accordingly transfer the brain mask and segmentation to the MRI image space. The toolbox package includes two home-made rat brain templates for 9.4 T and 14.1 T field strengths. These templates were generated from T_2_-weighted Turbo-Rapid acquisition with relaxation enhancement (Turbo-RARE) images (see Demonstration data acquisition section for details) and incorporate segmentations based on the SIGMA rat brain atlas [18]. Users also have the flexibility to use their own templates and segmentation schemes if desired.

##### Preprocessing

The MRSI data processing starts right after the MRI registration steps, and both are completely decorrelated (**Figure 2A**). The current version of the toolbox can handle different types of data available in the Bruker system, such as FID-MRSI, spin echo (SE)-MRSI, and point-resolved spectroscopy (PRESS)-MRSI, as long as they are reconstructed via the ParaVision software. First, metabolite MRSI data undergo water signal removal using the Hankel singular value decomposition (HSVD) method [19]. The resulting water-free metabolite data are then used for further processing steps, including the computation of a signal-to-noise ratio (SNR) map using the metabolite N-acetylaspartate (NAA). Subsequently, water MRSI data are processed to generate a linewidth map and a water power map. These SNR and linewidth maps serve as key metrics for quality control later in the processing pipeline. To exclude non-brain regions, both metabolite and water data are masked using the brain mask obtained from the co-registration step. Two optional preprocessing steps are provided in the *Processing Steps* panel: One is retrospective *Lipid suppression*, implemented based on singular value decomposition (SVD). This approach assumes that lipid and metabolite signals are orthogonal in the time or frequency domain and do not spatially overlap [20, 21]. The process begins by delineating brain and scalp regions using a mask of the water power map. SVD is then applied to the scalp voxels to construct an orthogonal basis representing lipid components. The rank of this basis is determined by assessing the energy ratio between brain and scalp regions after applying the corresponding projection operator as E_brain_/E_Skull_ ≥ α (default: α = 0.8). It is noted that in the toolbox, *Lipid suppression* only applies to metabolite data. Another method is backward linear prediction (called *Fillgaps* in the toolbox), implemented with an autoregressive algorithm (MATLAB’s *fillgaps* function) to reconstruct data points missing due to the inherent acquisition delay (AD) in FID-MRSI sequences. The number of missing points in FID is determined by the acquisition bandwidth (BW) and the AD time, and the prediction is applied to the FID signals [22]. *Fillgaps* applies to both metabolite and water data to ensure they are the same length in the quantification process. Once the processing is successfully performed, each spectrum corresponding to each voxel location in both metabolite and water data is saved into a *\*.RAW* file for LCModel fitting.

##### Fitting and Quantification

Fitting and quantification use the LCModel software [23] with a basis set containing spectral profiles of metabolites and macromolecules. The fitting process starts with the generation of \**.CONTROL* files containing data specification and all LCModel parameter configurations, some of which are specified in the *MRSI parameters* window (see the Practical guidelines section of the MRSI processing pipeline for details). Then, LCModel uses all information in the \**.CONTROL* file to handle the fitting and quantification. Upon successful processing, output files such as *\*.coord*, *\*.ps*, and *\*.table* are generated and stored in the results directory. It is necessary to note that although the methods presented in this work are primarily described for ^1^H MRSI data, the toolbox is capable of handling X-nuclei MRSI data. However, the pipeline must be carefully adapted for each specific nucleus, followed by thorough testing and validation.

#### 2.1.2. Practical guidelines

Using FID, SE demonstration datasets acquired at 9.4 T and 14.1 T, detailed in the Demonstration data acquisition section, the typical processing workflow is illustrated in **Figure 2B (**top-left) and comprises the following steps:

##### Step 1

Open the *MRSI Parameters* window by selecting the *MRSI Parameters* button (**Figure 2B**, top-right).

(1-1) Specify the directory containing the installed LCModel and select the appropriate metabolite basis set for the data type. For *Fillgaps* in particular, the basis set assuming no first-order phase effects (i.e., no AD) is needed to fit spectra reconstructed by the backward linear prediction. The metabolite basis sets are provided with demonstration datasets.
(1-2) Define LCModel parameters such as controlling baseline flexibility (DKNTMN, default: 0.25), prior phasing information such as DEGZER (default: 0), SDDEGZ (default: 999), DEGPPM (default: 0), SDDEGP (default: 0), and fitting ppm range specified in the configuration file (default: starting from 4.1 to 0.2 ppm), NRATIO (default: 0), WCONC (default: 44444), the internal reference metabolite (default: Cr+PCr), and its concentration (default: 8). Default parameters defined in the toolbox were used in this work, except for SDDEGP = 5 for 9.4 T and SDDEGP = 0 for 14.1 T datasets.
(1-3) Configure metabolite handling within the basis set (e.g., usage, combination, or omission).
(1-4) Save the configuration to a file. Note that using an existing filename will overwrite the previous configuration; assigning a new name prevents overwriting.
(1-5) Select template and atlas for anatomical co-registration. Template, brain mask, anatomical segmentation, and their labels for each field strength (9.4 T and 14.1 T) are included in the toolbox package.

##### Step 2

Specify the directories for data input and result output. It is important to assign unique study names (i.e., result folder names) for different analyses to prevent overwriting.

##### Step 3

Select processing options. There are two preprocessing options for two-dimensional (2D) or three-dimensional (3D) data: *Lipid suppression* and backward linear prediction (*Fillgaps*). If data are acquired from a 3D MRSI sequence, the *3D MRSI* checkbox and the number of slices should be specified.

##### Step 4

Input the data information, such as MRI, metabolite, and reference water MRSI scan numbers, slice number for brain regional anatomy, reconstruction number, the number of slices averaged for the display, and brain orientations if differences across scanners (e.g., *Head prone* or tail prone in the Bruker system). For demonstration data in this work, we used the center slice of the MRSI slab for brain regional anatomy, reconstruction number 2, which applies Hamming *k*-space filtering, and only selected *Head prone* for 9.4 T data.

##### Step 5

Run the analysis. Several message dialog boxes appear to inform the progress: Co-registration, HSVD, Linewidth, SNR, preprocessing steps such as *Lipid suppression* and *Fillgaps* if specified, and saving data to *\*.RAW* files for LCModel, quantification using LCModel, creation of metabolite map, etc. After completing the analysis, the relative concentration NAA map for the whole brain slice is the default display overlaid on the anatomical image.

##### Step 6

Explore display and output options (**Figure 2B**, top-left). The *Quality control* is to control the voxels that appear in the map using metabolite SNR (from NAA-based estimation or LCModel output), reference water linewidth, and Cramer-Rao lower bound (CRLB) percentage. The color map scaling is also set in this function. For demonstration purposes, we applied SNR ≥ 10, linewidth ≤ 1.25 times the averaged linewidth, and CRLB < 30%. A specific metabolite map for a specific brain region or whole brain slice can be optionally chosen by selecting one from the *Metabolite* and *Region* dropdowns. Brain mask, water power, SNR, linewidth, zero-order phase, and first-order phase maps are also listed in the *Metabolite* dropdown. Additionally, the CRLB map is also possible to display for quality assessment, as it is listed in the *Concentration* dropdown. The *Spectral display* window allows the user to inspect spectra at a chosen voxel. To do this, *Change voxel* must be enabled before selecting the desired voxel location (**Figure 2B**, bottom-left). This window also provides options to display spectra from spectroscopic *Data*, LCModel’s *Fit*, *Baseline,* and *Residuals*; the different quality metrics such as SNR, linewidth calculated during the processing, and by LCModel; and manual axis scaling if needed. The *Manual voxel selector* window is available for defining custom regions of interest, instead of using atlas-based regions for MRSI data masking (**Figure 2B**, bottom-left). Depending on the type of concentration (*absolute* or *relative*) specified in the *Concentration* dropdown, the *Concentration table* lists metabolite names, mean concentrations, standard deviations, and voxel counts for the chosen brain region that passed the quality control thresholds. The *Brain volumetry table* provides calculations of brain areas based on co-registered brain segmentation. Both concentration and brain volumetry tables can be saved as MATLAB data files in the result folder. The *Save map* allows users to store the currently displayed maps in the result folder. Additionally, if toolbox outputs, such as concentration tables, are saved in the result folder, it is possible to perform statistical tests using the *Statistical tool* (**Figure 2B**, top-left, and **Figure S1**). Options to select testing results, metabolite names, type of statistics, and their parameters [e.g., Student t-test, one/two-way analysis of variance (ANOVA), Bonferroni’s multiple comparisons, one/two-sided *p*-value], and to display the testing results (i.e., bar graph, result table) are all available.

To only view the results of the already processed/quantified data without reprocessing, first, all information, especially data and result directories, study name, data specifications, processing options, and parameter configuration, should be specified the same as when they were previously processed, then select the *MRSI analysis* button. Two or three message dialog boxes may appear, depending on the status of the toolbox. The message dialog box 1 asks if redoing image co-registration or keeping the preexisting one. The message dialog box 2 asks whether or not to restart the processing pipeline. The message dialog box 3 asks whether or not to reprocess the data. If the tool is newly launched (i.e., white in the metabolite map window), message dialog boxes 1 and 3 sequentially appear, and *Keep preexisting* and *No* should be respectively chosen. If the tool is in use (i.e., image or map shown in metabolite map window), message dialog boxes 1, 2, and 3 sequentially appear, and *Keep preexisting*, *Yes*, and *No* should be respectively chosen. The whole-slice NAA maps relative to total Creatine (tCr) concentration will appear as the default if the reloading process is successfully done.

### 2.2. SVS processing pipeline

#### 2.2.1. Processing methods

Vendor-reconstructed data (e.g., *fid/ser* files generated by Bruker ParaVision-360) are imported into the toolbox through the *Load SVS/DWS* interface (**Figure 3A**). If necessary, the data are automatically reformatted to ensure compatibility with subsequent processing steps. When multiple datasets are provided, each processing step is applied sequentially to every dataset before advancing to the next stage of the workflow. Raw data are then processed using the data processing module, which is adapted from the FID-A framework [10]. This module provides four optional preprocessing features: Image-selected in vivo spectroscopy (ISIS) handling, spectral alignment, outlier removal, and small voxel handling. Each feature, along with its associated parameters, can be configured and saved via the SVS/DWS *Parameter* window.

- ISIS handling: For datasets acquired using ISIS localization, the raw data are first separated into two subsets corresponding to odd and even shots within the ISIS scheme. Subsequent processing steps are applied independently to each subset.
- Spectral alignment: All acquisitions are aligned to their median spectrum using the NAA peak at 2.01 ppm as a reference. Adjustable parameters include line broadening (default: 12 Hz), frequency range limits, and maximum alignment time (default: 0.5 s).
- Outlier removal: When data exhibit excessive noise, this feature identifies and excludes outlier shots based on a rejection threshold (default: 1.5 SD) specified in the configuration file. This step can improve overall spectral quality before data combination.
- Small voxel handling: For ISIS data acquired from small voxels with low SNR, this function merges odd and even shots into a single dataset before further processing. Note that this reduces the total number of shots by half.

**Figure 3.**
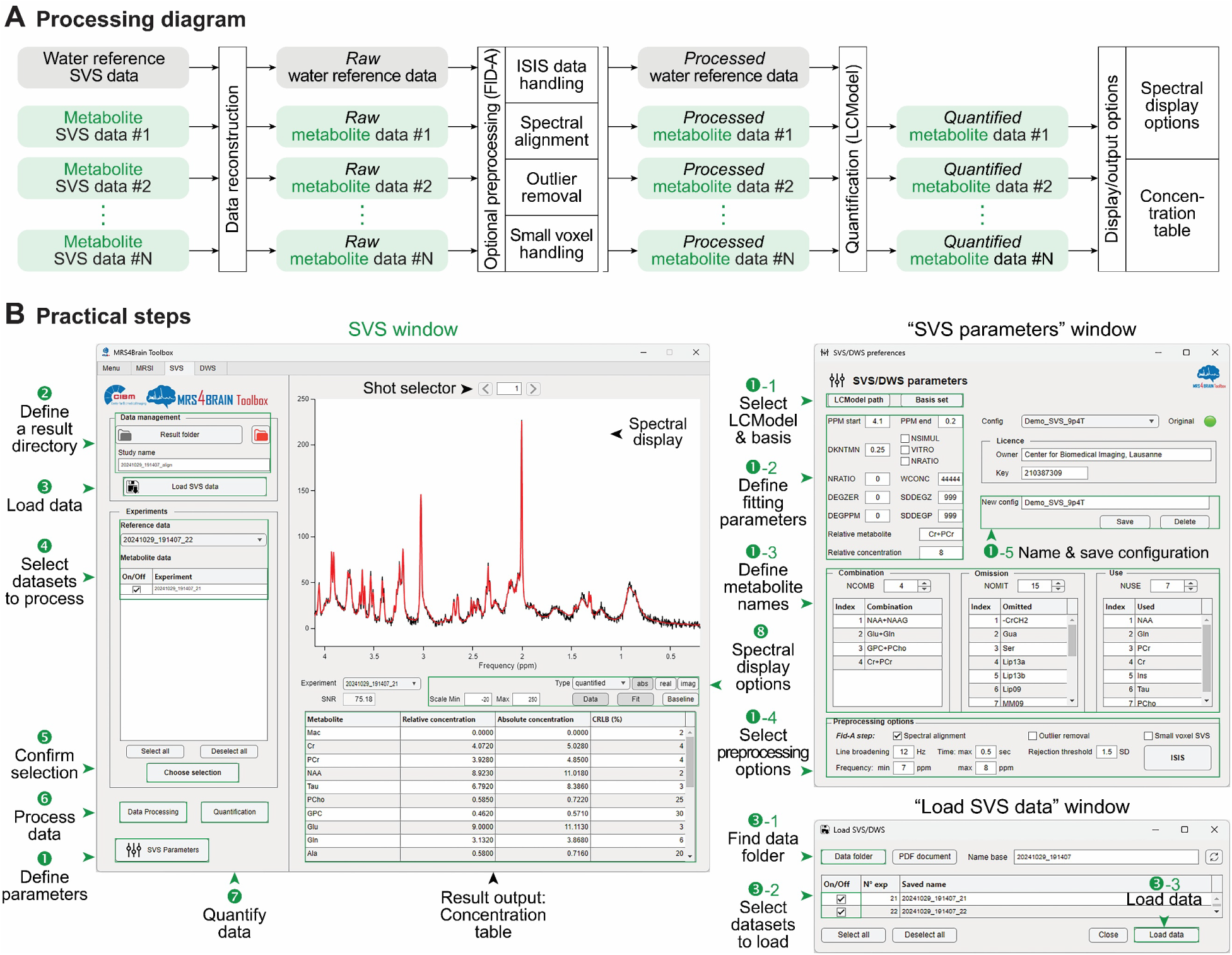
Processing pipeline and practical steps for SVS data. (**A**) Processing pipeline diagram from data reconstruction, processing to result display, and outputs (**B**) Typical workflow for processing an SVS dataset. Left: The main SVS window displays steps 1 to 8, guiding users through parameter configuration, data loading and processing, and selecting display and output options. Top-right: The *SVS/DWS Parameters* window (step 1) for defining parameter configurations. Bottom-right: The *Load SVS data* window (step 3), illustrating the procedure for loading datasets.

Following these steps, the preprocessed data are combined for quantification. LCModel fitting is then performed, with adjustable parameters for controlling baseline flexibility (DKNTMN, default: 0.25), prior phasing information such as DEGZER (default: 0), SDDEGZ (default: 999), DEGPPM (default: 0), SDDEGP (default: 999), and fitting ppm range specified in the configuration file (default: starting from 4.1 to 0.2 ppm), NRATIO (default: 0), WCONC (default: 44444), the internal reference metabolite (default: Cr+PCr), and its concentration (default: 8). The *\*.RAW* file is prepared as input for LCModel. Upon successful quantification, output files such as *\*.CONTROL*, *\*.coord*, *\*.ps*, and *\*.table* are generated in the results directory. The main SVS window subsequently displays individual fitted spectra and metabolite concentration values in the table.

#### 2.2.2. Practical guidelines

Using stimulated echo acquisition mode (STEAM) and spin echo full intensity acquired localized (SPECIAL) [24] SVS demonstration datasets acquired at 9.4 T and 14.1 T, detailed in the Demonstration data acquisition section, the typical processing steps are the following (**Figure 3B**, top-left):

##### Step 1

Open the *SVS/DWS Parameters* window by selecting the *SVS Parameters* button (**Figure 3B**, top-right).

(1-1) Specify the directory containing the installed LCModel and select the appropriate metabolite basis set for the data type. The metabolite basis sets are provided with demonstration datasets.
(1-2) Define LCModel parameters (e.g., DKNTMN, DEGZER, SDDEGZ, DEGPPM, SDDEGP, fitting frequency range, etc.), the internal reference metabolite, and its concentration. Default parameters defined in the toolbox were used in this work (see Processing methods section of SVS processing pipeline for default values).
(1-3) Configure metabolite handling within the basis set (e.g., usage, combination, or omission).
(1-4) Select preprocessing options. For the STEAM data, we applied spectral alignment with outlier removal. Small voxel and ISIS options are not selected. For the SPECIAL data, we applied ISIS handling, small voxel handling, spectral alignment, and outlier removal. All other parameters remain at default values.
(1-5) Save the configuration to a file. Note that using an existing filename will overwrite the previous configuration; assigning a new name prevents overwriting.

##### Step 2

Specify the directory for output. It is important to assign unique study names (i.e., result folder names) for different analyses to prevent overwriting.

##### Step 3

Open the *Load SVS/DWS data* window by selecting the *Load SVS data* button (**Figure 3B**, bottom-right).

(1-1) Specify the data directory.
(1-2) Select all datasets used for analysis. Options to open an experiment note PDF file and rename data prefixes are available if necessary.
(1-3) Complete the process by selecting the *Load data* button.

##### Step 4

Select metabolite datasets for analysis and a water dataset for reference.

##### Step 5

Confirm selection by selecting the *Choose selection* button. If successfully processed, the raw spectrum (Type: *raw*) will appear in the *Spectral display* window. It is possible to inspect the individual shot spectra using the shot selector at the top of the *Spectral display* window.

##### Step 6

Perform preprocessing steps by selecting the *Data processing* button. If successfully processed, a combined spectrum (Type: *processed+sum*) will appear in the *Spectral display* window. It is possible to review the processed data before combination by selecting *processed* in the *Type* dropdown and using the shot selector.

##### Step 7

Perform quantification by selecting the *Quantification* button. If successfully quantified, fitted spectra (Type: *quantified*) will display alongside data in the *Spectral display* window.

##### Step 8

Options to visualize and output results. Spectra could be displayed in absolute (*abs*), real (*real*), or imaginary (*imag*) formats for raw, processed, and combined data. For quantified data, additional options such as *Data*, *Fit*, and *Baseline* are available. Results, including spectra and concentration tables from different datasets, could be viewed by selecting the corresponding dataset name in the *Experiment* dropdown. All data are stored in the folder named in step 2, which contains subfolders as *raw, processed,* and *quantified,* corresponding to steps in the pipeline. If *Outlier removal* is used, text files containing removed shot numbers are also saved in the folder.

To only view the results of the already processed/quantified data without reprocessing, first, it is necessary to specify the data and result directories. Once the study name is entered correctly, the toolbox will automatically reload the existing results. When the fitted spectrum is overlaid with the SVS data in the Spectral window, it confirms that the results have been successfully reloaded.

### 2.3. DWS processing pipeline

#### 2.3.1. Processing methods

The DWS data processing pipeline (**Figure 4A**) is adapted from the work described by Jessie et al. [25] and closely follows the structure of the SVS framework. It employs the same *Load SVS/DWS* interface for data import and the *SVS/DWS Parameters* window for defining processing parameters. Preprocessing capabilities include ISIS handling, spectral alignment, and outlier removal, followed by quantification using LCModel.

**Figure 4.**
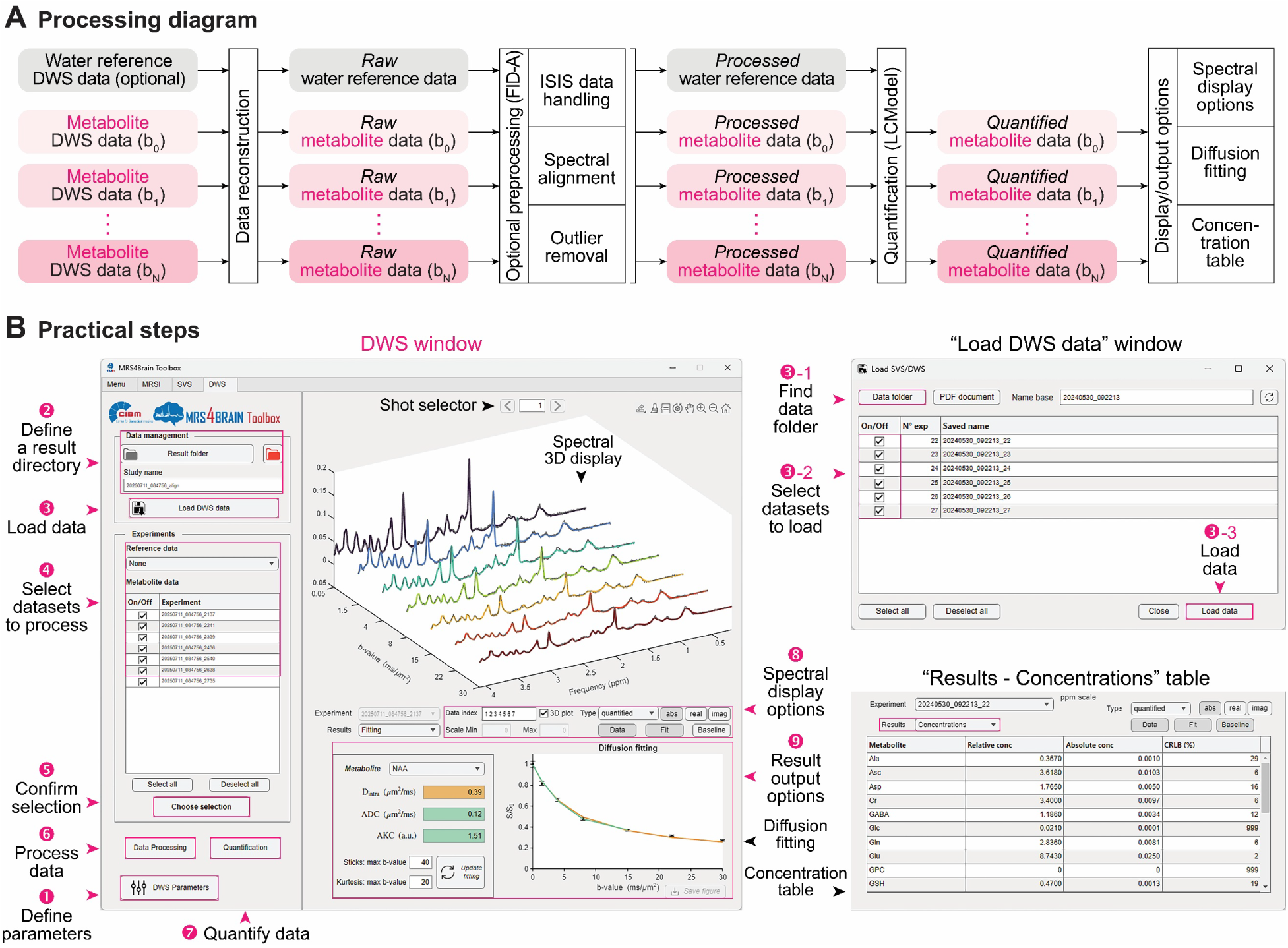
Processing pipeline and practical steps for DWS data. (**A**) Processing pipeline diagram from data reconstruction, processing to result display, and outputs (**B**) Typical workflow for processing a DWS dataset. Left: The main DWS window shows steps 1 to 9, guiding users through parameter configuration, data loading and processing, and selecting display and output options. Top-right: The *Load DWS data* window (step 3) demonstrates the procedure for loading datasets. Bottom-right: The “concentration” table as an output option, alongside diffusion fitting.

For spectral visualization, in addition to displaying individual spectra, the pipeline supports 3D plotting of all DWS-fitted spectra across different b-values, enabling assessment of signal attenuation as a function of b-value for a given diffusion time. Similar to the SVS workflow, a “concentration” table is provided as one of the output options. If the reference dataset, such as water data, is not specified, this table contains values in arbitrary units. An additional feature is diffusion fitting, which estimates diffusion properties for each metabolite. This functionality is implemented using a randomly oriented stick [26] and kurtosis [27] models. It should be noted that the b-values displayed in the toolbox are extracted directly from the scan parameters specified in the method file. Corrections for factors such as cross-term interactions must be estimated externally to obtain accurate final results [28].

#### 2.3.2. Practical guidelines

Using diffusion-weighted SPECIAL (dSPECIAL) [25] DWS demonstration datasets acquired at 9.4 T and 14.1 T, detailed in the Demo data section, the typical processing steps are the following (**Figure 4B**, top-left):

**Steps 1 to 7** are closely aligned with steps 1 - 7 described in the practical guidelines for SVS data (**Figure 4B**, top-left and top-right). For preprocessing options in step 1, we selected ISIS handling, spectral alignment, and outlier removal. All other parameters remain at default values.

##### Step 8

Options to visualize results. In addition to step 8 described in the practical guidelines for SVS data, 3D plotting is available in the DWS processing pipeline to visualize signal attenuation across b-values. When the *3D plot* option is selected, all spectra are displayed in 3D, with options to show *Data*, *Fit*, and *Baseline* also supported in this view. Datasets are automatically sorted in ascending order by name; however, if data acquisitions at different b-values were performed in random order, they can be manually rearranged in ascending b-value order using the *Data Index* box for display. Unlike SVS, DWS spectra, particularly at high b-values, exhibit highly reduced SNR due to the influence of diffusion gradients or motions. Therefore, it is essential to evaluate individual shots at each processing step using the shot selector.

##### Step 9

Options to output the results. Similar to the SVS processing pipeline, the “concentration” table displays both relative and absolute concentrations, along with CRLB values for each metabolite (**Figure 4B**, bottom-right). The diffusion fitting feature offers additional tools to examine diffusion properties of each metabolite in the Metabolite dropdown, either through intra-stick diffusivity (D_intra_), apparent diffusion coefficient (ADC), and apparent kurtosis coefficients (AKC), or by viewing fitted curves. It is possible to restrict the maximum b-values for diffusion fitting models using the corresponding input fields and apply changes by clicking the *Update fitting* button. To assist this, the CRLB values in the table can help determine an appropriate range of b-values or identify which metabolites are accurately fitted.

### 2.4. Demonstration data acquisition

Wistar adult rats (Charles River Laboratories, L’Arbresle, France) under 1.5 to 2.5% isoflurane anesthesia were used for in vivo acquisitions. The protocol was approved by the Committee on Animal Experimentation in the Canton of Vaud, Switzerland. The body temperature of the animals was kept at 37.5 ± 1.0 °C by circulating warm water and measured with a rectal thermosensor. The respiration rate and body temperature were monitored using a small-animal monitor system (SA Instruments, New York, USA). During data acquisitions for MRI, MRSI, SVS, and DWS, all animals were placed in an in-house-built or in a Bruker-manufactured holder, with their head fixed in a stereotaxic system using a bite bar and a pair of ear bars.

Each type of demo data was acquired at two field strengths (9.4 T and 14.1 T). The horizontal 9.4 T magnet (Magnex Scientific, Yarnton, UK) has a 660 mT/m peak strength and 4570 T/m/s slew rate shielded gradient set (Bruker B-GA12S HP) interfaced to a Bruker console (BioSpec AVANCE NEO, ParaVision 360 v3.5), and using a quadrature volume transmit coil and a four-channel cryogenic receive coil specialized for rat head (CryoProbe, Bruker), whereas the horizontal 14.1 T magnet (Magnex Scientific, Yarnton, UK) has a 1 T/m peak strength and 5500 T/m/s slew rate shielded gradient set (Resonance Research, Billerica, USA) interfaced to a Bruker console (BioSpec AVANCE NEO, ParaVision 360 v3.3), and using a home-made transceiver quadrature surface coil (two 1.8 × 1.6 cm loops, covering a curved 1.8 × 2.7 cm^2^ surface with 14 mm radius of curvature).

For reference, coronal and axial MRI images were acquired on both systems using a T_2_-weighted Turbo-RARE image. Scan parameters were repetition time (TR) = 4100 ms, echo time (TE) = 27 ms, number of averages = 10, RARE factor = 6, 128 × 128 matrix, field of view (FOV) = 24 × 24 mm, slice thickness = 0.3 mm (9.4 T) or 0.2 mm (14.1 T). The shimming procedure described by Simicic and Alves et al. [6] and demonstrated by Cudalbu and Alves [29, 30] was used.

#### 2.4.1. MRSI data

##### FID-MRSI

Metabolite data were acquired from a single slice, mainly covering the hippocampus and striatum. FID-MRSI sequence was used at 9.4 T with the following parameters: TR = 822 ms (9.4 T) or 812 ms (14.1 T), AD = 1.3 ms, flip angle (FA) = 55° (9.4 T) or 52° (14.1 T), BW = 5000 Hz (9.4 T) or 7143 Hz (14.1 T), 768 (9.4 T) or 1024 (14.1 T) spectral points, 31 × 31 matrix, FOV = 24 × 24 mm, voxel size = 0.77 × 0.77 × 2 mm^3^, single average, water suppression using variable power radiofrequency pulses and optimized relaxation delays (VAPOR) scheme, lipid suppression using 12 (9.4 T) or six (14.1 T) saturation bands placed on cranial regions. Water data were acquired using the same parameters, except that water suppression was disabled.

##### SE-MRSI

Metabolite and water data were acquired from the same experimental setup procedure and scanning parameters as used in FID-MRSI at 9.4 T, except that the SE-MRSI sequence was used at 9.4 T with TR = 2000 ms, TE = 3.3 ms, FA = 90°/180°.

More detailed methods and scanning parameters for MRSI data can be found in our other studies [6, 31].

#### 2.4.2. SVS data

##### STEAM SVS

Metabolite data were acquired from a 2.8 × 2 × 2 mm^3^ voxel in the hippocampus. STEAM sequence was used with the following parameters: TR = 4000 ms, TE = 3 ms, TM = 10 ms, BW = 5000 Hz, 4096 spectral points, 16 averages, and 16 repetitions, water suppression using VAPOR. Water data were acquired using the same parameters, except that water suppression was disabled.

##### SPECIAL SVS

Metabolite data were acquired from a 2.8 × 2 × 2 mm^3^ voxel in the hippocampus. SPECIAL sequence [24] was used at 14.1 T with the following parameters: TR = 4000 ms, TE = 9.3 ms, BW = 7143 Hz, 4096 spectral points, 256 repetitions, water suppression using VAPOR, and an additional water suppression pulse inserted in the mixing time (TM). Water data were acquired using the same parameters, except that water suppression was disabled.

#### 2.4.3. DWS data

Metabolite data were acquired from a 10 × 2.5 × 4 mm^3^ (9.4 T) or 7 × 5 × 5 mm^3^ (14.1 T) voxel in the hippocampus. dSPECIAL sequence [25] was used at 9.4 T or 14.1 T with the following parameters: TR = 3000 ms, TE = 9.5 ms, TM = 150 ms (9.4 T) or 40 ms (14.1 T), BW = 5000 Hz (9.4 T) or 7143 Hz (14.1 T), 4096 spectral points, b-value = 0.05, 1.5, 4, 8, 15, 22, 30 ms/µm^2^ with 64, 64, 96, 96, 128, 128, 192 repetitions respectively (9.4 T) or 0.05, 1, 3, 5, 10, 20 ms/µm^2^ with 160, 160, 160, 160, 320, 320 repetitions respectively (14.1 T), water suppression using VAPOR and an additional water suppression pulse in TM.

#### 2.4.4. Basis set

Basis sets were generated using metabolites simulated by NMRScopeB from jMRUI [32–34], employing the same acquisition sequence and parameters as the corresponding in vivo measurements and the experimentally measured macromolecule signal under the same conditions. Simulations incorporated published J-coupling constants and chemical shifts [35–37]. Each basis set was tailored to the specific scan configuration of the dataset, including sequence type, TE, AD, magnetic field strength, and other relevant parameters.

## 3. Results

### 3.1. MRSI data from different sequences and field strengths

Following the described practical guidelines, we first demonstrate the MRSI functionalities using the options available in the MRSI processing pipeline. There are three datasets from different sequences and field strengths used for this purpose: FID-MRSI data at 9.4 T (**Figures 5A** to **5D**), SE-MRSI data at 9.4 T (**Figures 5E** and **5F**), and FID-MRSI data at 14.1 T (**Figures 5G** and **5H**). For FID-MRSI, **Figure 5A** shows brain mask edges overlaid on the anatomical image, obtained via co-registration between the acquired MRI image and the anatomical template (**Figure 5A**, top). Similarly, the linewidth map (**Figure 5A**, middle) estimated from water MRSI data and SNR maps calculated using metabolite MRSI data (more specifically NAA) after applying the brain mask (**Figure 5A**, bottom) are shown as outputs from the toolbox. With the default processing option, we obtained NAA and myo-inositol (Ins) metabolite maps for the whole brain slice, and with regional highlights such as the striatum delineated by brain segmentations. In addition, CRLB maps extracted from LCModel outputs were included to assess the fitting accuracy, as illustrated in the first six columns of **Figure 5B**. These results from the metabolite map window were simply exported using the *Save map* button. As another output of the toolbox, the spectrum for a voxel selected in the *Spectral display* window is presented in the last column of **Figure 5B**. The result can also be quantitatively presented using the *Concentration table* (**Supplementary Table 1**) and, in this case, tCr as internal reference, displaying averaged values from counted voxels in the hippocampus. To demonstrate the capability of the preprocessing options, we repeated the analyses, but processing with *Lipid suppression* (**Figure 5C**) and with *Fillgaps* (**Figure 5D**). Slight changes in the maps upon applying the preprocessing steps were observed (**Figures 5C** and **5D**). In results from processing with *Fillgaps*, the output provides fully phased spectra, indicating the effects of backward linear prediction of missing points in FID (see the last column of **Figure 5D**).

**Figure 5.**
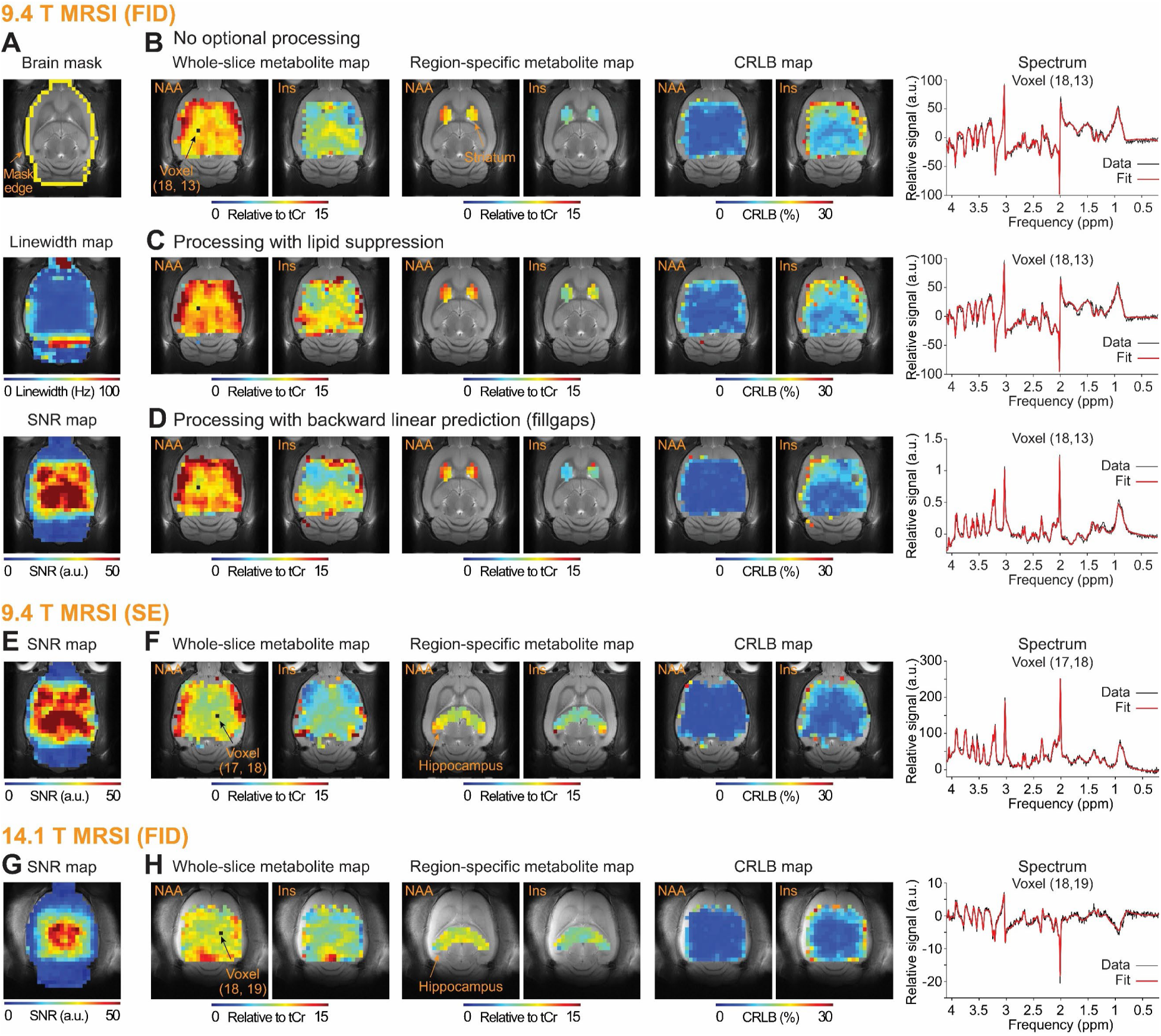
Results from processing MRSI demonstration datasets. (**A** to **D**) FID-MRSI data at 9.4 T. (A) Top to bottom: brain mask output from the co-registration step, linewidth map from water data, and SNR map from metabolite data after applying the brain mask. (B) Processing with default options. Left to right: whole-slice and region-specific (striatum) metabolite maps, CRLB maps for NAA and Ins, and fitted spectrum (red) overlaid on data from a representative voxel (black). (C) The same as (B), but for processing data with an additional lipid suppression option. (D) The same as (B), but for processing data with an additional backward linear prediction (fillgaps) option. (**E** and **F**) SE-MRSI data at 9.4 T. (E) SNR map from metabolite data after applying the brain mask. (F) Processing with default options. Left to right: whole-slice and region-specific (hippocampus) metabolite maps, CRLB maps for NAA and Ins, fitted spectrum (red) overlaid on data from a representative voxel (black). (**G** and **H**) The same as (E and F), but for FID-MRSI data at 14.1 T. Voxels in the metabolite maps passed the quality control thresholds as SNR ≥ 10, linewidth ≤ 1.25 times the averaged linewidth, and CRLB < 30%. Metabolite maps are presented relative to tCr fixed at 8 mmol/kgww. Black dots in the NAA map indicate voxel locations for the representative spectra in the last column.

For other sequences, the toolbox produced excellent maps and spectra, as shown in **Figures 5E** and **5F** for SE-MRSI at 9.4 T, and **Figures 5G** and **5H** for FID-MRSI at 14.1 T, demonstrating the capability of the *MRS4Brain Toolbox* to process MRSI data across a broad range of dataset types.

### 3.2. SVS data from different sequences and field strengths

Next, we examined the SVS functionalities using the available options in the SVS processing pipeline. Two datasets acquired in the hippocampus using different sequences and field strengths were examined: STEAM data at 9.4 T (**Figures 6A** to **6D**) and SPECIAL data at 14.1 T (**Figures 6E** and **6F**). For the STEAM dataset, selecting the *Choose Selection* option displayed the raw spectra, allowing users to inspect individual spectra from the first to the last shot (**Figure 6A**, top and bottom, respectively). Upon selecting the *Data processing* button, the pipeline generated both *processed* spectra (**Figure 6B**) and *combined* spectra (**Figure 6C**). By default, the *combined* spectrum, including *Spectral alignment* and *Outlier removal*, was exhibited. *Processed* spectra for each shot (**Figure 6B**) could be viewed by selecting *processed* in the *Type* dropdown. Following LCModel fitting, the *quantified* spectrum was shown with the measured data in black and the fitted one in red (**Figure 6D**). Also, a results table was generated, listing metabolite names, concentrations relative to tCr, absolute concentrations, and CRLB values (**Supplemental Table 2**). Processing of the SPECIAL dataset followed the same general procedure as used in STEAM data analysis. However, because the data were acquired using an ISIS localization scheme, ISIS handling was required, in addition to Spectral Alignment and Outlier Removal. Small voxel handling was also enabled for this demonstration (**Figure 6E**). The final output showed the fitted LCModel spectrum overlaid on the measured data (**Figure 6F**), confirming successful analysis of the SPECIAL dataset.

**Figure 6.**
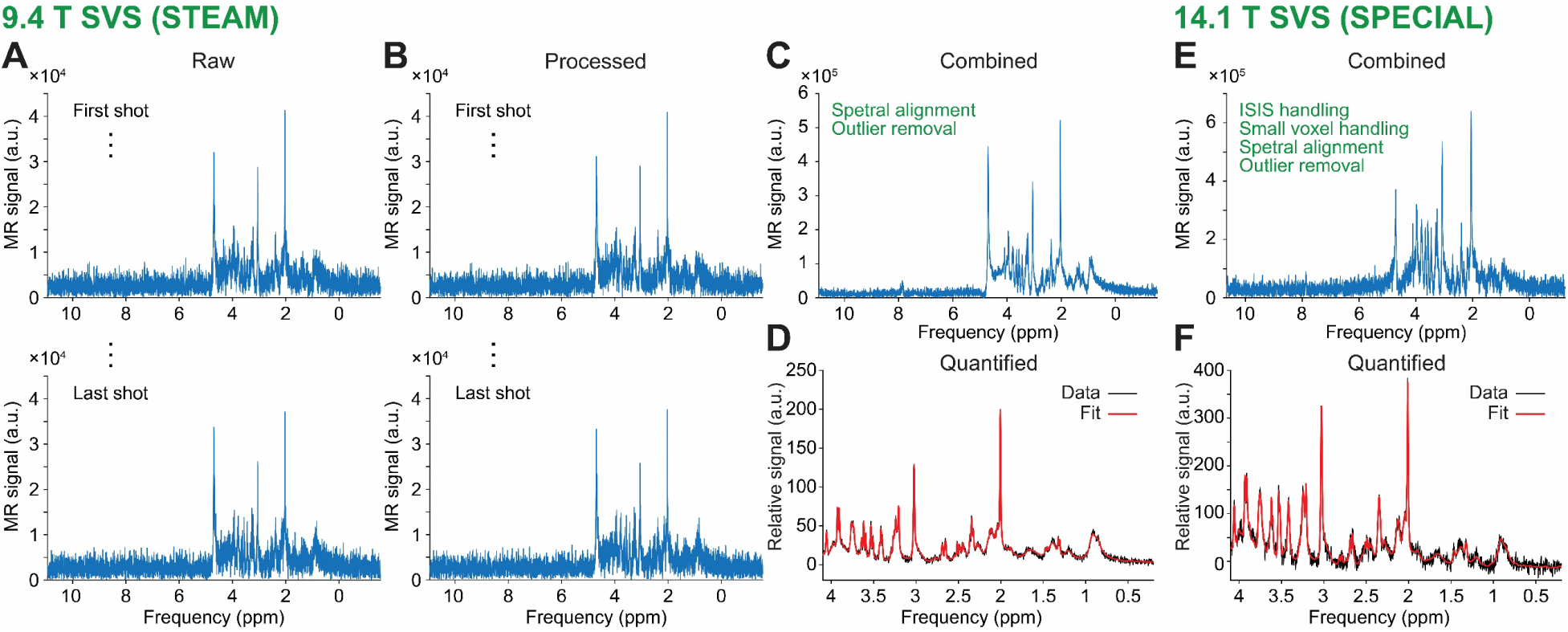
Results from processing SVS demonstration datasets. (**A** to **D**) STEAM SVS data at 9.4 T. (A) Unprocessed (raw) spectrum displayed individually from the first shot (top) to the last one (bottom), after selecting *Choose selection*. (B) The same as (A), but after applying spectral alignment. (C) Spectrum with all shots combined, excluding outliers. (D) LCModel’s fitted spectrum (red) displayed with data (black). (**E** and **F**) SPECIAL data at 14.1 T. The same as (C and D) but processed with additional preprocessing steps such as small voxel and ISIS handling.

Overall, these results demonstrate that the SVS processing pipeline can reliably process SVS datasets acquired with different sequences and at different field strengths using the full suite of functionalities available in the toolbox.

### 3.3. DWS data at different field strengths

Finally, we evaluated the DWS functionalities using the available options in the DWS processing pipeline. Two datasets acquired in the hippocampus with the dSPECIAL sequence at different field strengths were analyzed: 9.4 T (**Figures 7A** to **7C**) and 14.1 T (**Figures 7D** to **7F**). Similar to the SVS processing pipeline, spectra for each average/shot, either *raw* or *processed*, are displayed in the *Spectral window* once each processing step is completed. Because the dSPECIAL data were acquired using an ISIS localization scheme, *ISIS handling* was required in addition to *Spectral alignment* and *Outlier removal*. After preprocessing, *combined* spectra for each b-value were generated (**Figures 7A** and **7D**). Selecting a dataset from the *Experiment* dropdown allows users to inspect the spectrum for a given b-value and assess preprocessing effects. Quantification is performed by selecting the *Quantification* button, which generates a fitted spectrum overlaid with the processed data, consistent with the SVS pipeline. Unique to the DWS pipeline, the *3D plot* (**Figures 7B** and **7E**) provides an efficient overview of all spectra within a single frame. This enables a quick evaluation of diffusion-related signal attenuation across b-values, even when datasets across b-values were acquired in random order, as data indices can be rearranged using the *Data index* box. In addition to a table summarizing concentration or arbitrary unit values (**Supplementary Table 3**), the toolbox offers *Diffusion fitting*, which displays the relative signal, S/S₀, computed as the signal at a given b-value normalized to the signal at the first b-value. In this demonstration, DWS data were fitted using either an empirical model (e.g., kurtosis) or a simplified biophysical model (e.g., the commonly used randomly oriented stick model) (**Figures 7C** and **7F**). Estimated parameters, including D_intra_, ADC, and AKC, are displayed as shown in **Figure 4B** (left). Because each model is valid over a specific b-value range, the toolbox allows users to select an appropriate fitting range. For the kurtosis model, suitable b-values typically extend up to 10 - 15 ms/µm^2^, depending on the metabolite [38].

**Figure 7.**
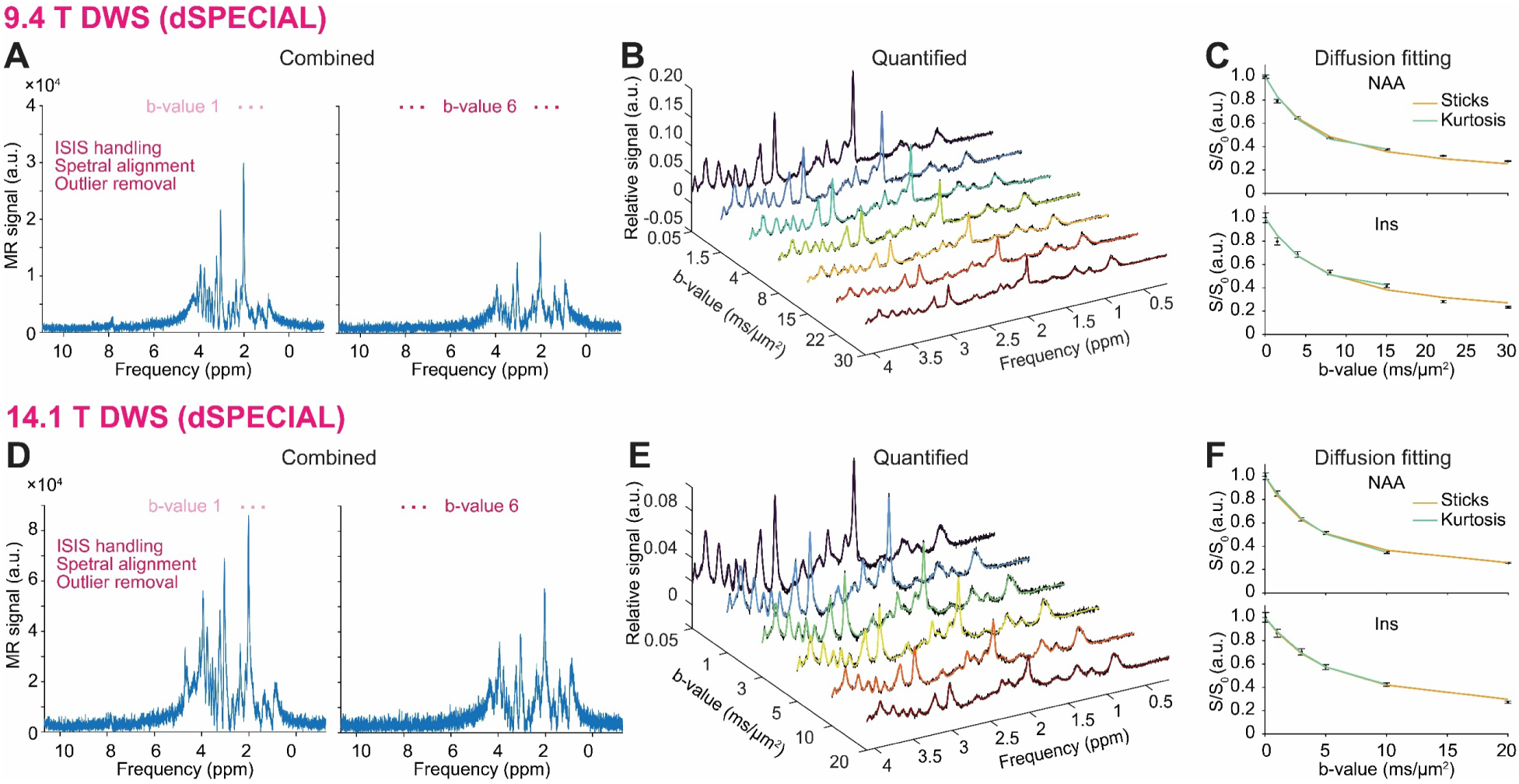
Results from processing DWS demonstration datasets. (**A** to **C**) dSPECIAL DWS data at 9.4 T. (A) Spectrum obtained by combining all shots, following processings using ISIS handling, spectral alignment, and outlier removal, for individual datasets: b-value 1 (left) and b-value 6 (right). (B) Spectra across all b-values displayed in 3D plotting mode, with LCModel’s fitted spectra (rainbow colors) overlaid on the data (black). (C) Diffusion fits using randomly oriented sticks (yellow) and kurtosis (green) models for NAA (top) and Ins (bottom), plotted as a function of b-values (**D** to **F**) The same as (A to C), but for dSPECIAL DWS data at 14.1 T.

Together, these results demonstrate that the DWS processing pipeline can robustly process various types of DWS datasets (e.g., dSPECIAL acquired at different field strengths) using the full set of functionalities implemented in the toolbox.

## 4. Discussion

We developed and presented herein the *MRS4Brain Toolbox*, a freely available software to streamline the processing of preclinical MRSI, MRS, and DWS datasets, even for non-spectroscopy experts. This comprehensive tool combines processing pipelines to process datasets acquired on preclinical Bruker scanners. For MRSI, it enables automated processing with semi-automatic quality control, removal of spurious signals, spectral fitting, brain segmentation, visualization, and co-registration of the metabolic maps, and built-in statistical tools. This allows streamlined processing of the huge amount of data generated in MRSI acquisitions, which would not be feasible manually in a reliable manner. Covering a comprehensive set of spectroscopy methods, it also includes processing pipelines for preprocessing and quantification of single-voxel spectroscopy datasets, as well as fitting of diffusion-weighted spectroscopy, allowing for a variety of uses for preclinical spectroscopic studies.

In the current implementation, the *MRS4Brain Toolbox* is compatible with datasets acquired on Bruker systems running ParaVision 360, which have been installed since 2019. As such, usage of the toolbox is limited to the data format predefined by the ParaVision software. One possibility to explore is the ability to read data in a MATLAB format, which would require the user to have their own data reading code to translate from any vendor format to a readable MATLAB file that could skip the reading step of the process. Additionally, while the toolbox manages to cover a variety of sequences, there exist some blind spots for advanced MRSI sequences, such as spatial-spectral encoding MRSI. Indeed, as the *MRS4Brain Toolbox* is designed to work with vendor-provided sequences to facilitate its use on many sites without having to program or install custom acquisition sequences, it only supports datasets that are already spatially reconstructed (in image space). To cover this issue, we plan to use the *rawdata.job0* file provided by ParaVision 360 containing the raw *k*-space data acquired, and to proceed to the sequence-specific reconstruction to fall back into a data structure more akin to what is found in *\*.fid* or *\*.fid_proc.64.* This future work may be implemented for both fully sampled and undersampled MRSI datasets. The undersampled dataset may further benefit from advanced compressed sensing reconstruction approaches such as low-rank total generalized variation (LR-TGV) [7, 39] or SPIRiT [40].

We have recently shown promising results using different denoising strategies, e.g., Marchenko-Pastur principal component analysis (MP-PCA), total variation (TV) regularization [7, 41, 42], stacked partial separability and spectral and temporal integration for SVD (SPIN-SVD), and tensor MP-PCA (tMPPCA) [43, 44]. Next versions of the MRS4Brain toolbox will integrate these denoising strategies to enable robust, automated workflows for diverse preclinical spectroscopy applications, as well as include thorough testing and validation of X-nuclei MRSI and MRS data sets combined with advanced metabolic modelling.

The implementation of advanced sequences can be extended to DWS since most sequences related to this modality used in research are custom-built. As it stands, the toolbox is optimized for sequences using a single b-value per acquisition. The toolbox may require adaptation for specific sequence implementations, such as those employing interleaved b-values and/or gradient directions within a single acquisition protocol [45, 46]. Including a monoexponential model would be useful for datasets with only a few small b-values, or including a rapid quality check to verify data consistency, consisting of spectra displays for each processing step. Beyond this, incorporating more sophisticated models would be highly beneficial for further applications and could help democratize the adoption of DWS by enabling the estimation of microstructural parameters such as soma radius and projection radius, as in the soma and neurite density imaging (SANDI) model [47]. These advanced models are particularly relevant because they allow moving beyond simple diffusion metrics to infer biologically meaningful features of tissue microstructure, such as cell body size and neurite geometry, which are critical for understanding brain organization and pathology. Models such as sphere-and-stick or cylinder could also be integrated to broaden the toolbox’s modeling capabilities and support more detailed microstructural analysis. Finally, processing and analysis that include multiple diffusion times would also be in line with recent publications [48, 49], offering more potential applications for DWS in both preclinical research. In upcoming versions of the toolbox, additional functionality will likely include advanced visualization tools for data processing steps. This would be particularly important to align with best practices in DWS [28], which require extensive quality checks since even minor animal movements can significantly affect the DWS signal. For example, useful visualization features could include overlaying individual repetitions to verify alignment and tools for outlier detection and removal, with an option to manually select specific outliers for exclusion.

Efficient brain function depends on the seamless integration of localized neural processing with long-range network connectivity, in which focal activations support specialized functions and long-range connections facilitate efficient information exchange between distributed brain regions. Together, these processes underpin normal and efficient cognitive functioning, sensory processing, motor control, and other behavioral operations. In light of this network-based perspective, the long-term trajectory of preclinical MRSI research is expected to move beyond characterizing region-specific alterations but toward characterizing the brain as an integrated network to better capture the diverse pathological processes associated with neurological disease [50, 51]. To support this shift, one of our aims is to extend the current analysis toolbox with a new feature capable of generating metabolic similarity matrices, enabling the evaluation of metabolic relationships across brain regions and advancing the study of network-level metabolic dysfunction.

The same atlas-based architecture that supports metabolic maps and connectivity analyses in MRSI can be extended to positron emission tomography (PET) studies, providing a natural multimodal bridge toward PET-derived metabolic flux maps, as can be obtained, for example, with Fluorodeoxyglucose (FDG)-PET [52], or metabolic connectivity in future implementations. PET represents a mature molecular imaging modality capable of quantifying pathway-specific metabolism and receptor-level binding affinity with high sensitivity, and its integration constitutes a logical expansion of the toolbox toward broader multimodal characterization of brain healthy function and pathologies. Recent work in rodent imaging has shown that reproducible tracer-derived biomarkers require MRI-based co-registration and atlas-defined regional sampling to ensure anatomical precision and reduce methodological variability [53]. In PET images, anatomical reference points are typically missing, with the PET intensity distribution reflecting tracer-specific molecular fate rather than underlying tissue structure, and some tracers show no uptake in major brain regions. Multimodal PET-computed tomography (CT) has become a standard multimodal approach to obtain fast anatomical reference images. However, PET-MRI offers superior soft tissue contrast, in particular in the brain, with better tissue differentiation and without the use of additional ionizing radiation. Moreover, the use of MRI equipment opens the way to all MR-based approaches, such as functional MRI for functional studies and MRSI for metabolic imaging. Within this framework, PET and MRSI provide complementary readouts: PET quantifies molecular kinetics (e.g., receptor binding or glucose utilization flux with FDG-PET, whereas MRSI reflects the intracellular biochemical state, its integrity, or perturbation in specific brain diseases. These coordinated measurements have shown their potential in rodent disease models, such as bile duct ligated (BDL) rats for the study of hepatic encephalopathy, where regional reductions in glucose metabolism, measured with FDG-PET, were shown to co-occur with perturbations in glutamatergic and osmotic metabolites detected by ^1^H-MRS [52]. Beyond such biological complementarity, recent methodological developments have demonstrated that dynamic FDG-PET can also be used to derive within-individual metabolic connectivity networks [54], providing metabolic information beyond static regional measurements. Together, these findings highlight that both MRSI and single-voxel spectroscopy can provide metabolic information that meaningfully complements PET-derived measures and therefore represent valuable approaches for future multimodal PET–MR integration. Future releases of the *MRS4Brain Toolbox* will incorporate PET processing by leveraging the existing ANTs-based registration, atlas segmentation, and regional statistical framework, enabling PET-MRI-atlas alignment and spatially resolved multimodal analyses with MRSI/MRS-derived metabolic profiles.

## 5. Conclusion

We introduced the *MRS4Brain Toolbox*, a freely available and comprehensive software designed to streamline the processing of preclinical MRS, DWS, and MRSI datasets, even for non-experts in spectroscopy. By integrating automated and semi-automatic pipelines for preclinical acquisitions such as Bruker data, the toolbox enables efficient handling of large-scale MRSI data through robust quality control, different processing steps, spectral fitting, brain segmentation, visualization, metabolic maps, and statistical analysis. Support for both SVS and DWS, including modelling, further extends its capability across a wide range of preclinical studies. By reducing manual workload and ensuring reproducibility, *MRS4Brain Toolbox* facilitates advanced preclinical spectroscopic research and promotes broader adoption of MRSI, MRS, and DWS methodologies in neuroscience.

## Data availability

The *MRS4Brain Toolbox* v.1.0 and datasets for demonstration purposes are publicly available on the GitHub repository: https://github.com/AlvBrayan/MRS4Brain-toolbox and https://www.epfl.ch/labs/mrs4brain/ressources/mrs4brain-toolbox/.

## Acknowledgements

We acknowledge the CIBM Center for Biomedical Imaging of the Ecole Polytechnique Fédérale de Lausanne (EPFL), the Université de Lausanne (UNIL), Université de Genève (UNIGE), the Hôpitaux Universitaires de Genève (HUG), and the Centre Hospitalier Universitaire Vaudois (CHUV) for providing expertise and resources to conduct this study. We thank Dr. Katarzyna Pierzchala for her support in generating segmentation labels for the rat brain templates, and Drs. Antoine Klauser and Dunja Simicic for valuable discussions during the development of the toolbox. We also thank Estelle Gerossier, Jocelyn Grosse, and Stefanita-Octavian Mitrea for their work in handling the animal experiments for the demonstration data acquisitions.

## Author contributions

Study design: Brayan Alves, Tan Toi Phan, Guillaume Briand, Cristina Cudalbu, Thanh Phong Lê; Conceptualization and programming: Brayan Alves, Tan Toi Phan, Guillaume Briand, Jessie Mosso, Jamie Near.

Data acquisition: Brayan Alves, Tan Toi Phan, Alessio Siviglia, Gianna Nossa, Jessie Mosso, Cristina Cudalbu.

Formal analysis and investigation: Brayan Alves, Tan Toi Phan, Alessio Siviglia, Gianna Nossa, Thi Ngoc Anh Dinh, Thanh Phong Lê, Cristina Cudalbu.

Writing - original draft preparation: Brayan Alves, Tan Toi Phan, Alessio Siviglia, Gianna Nossa, Eloise Mougel, Thi Ngoc Anh Dinh, Omar Zenteno, Bernard Lanz, Thanh Phong Lê, Cristina Cudalbu.

Writing - review and editing: Brayan Alves, Tan Toi Phan, Guillaume Briand, Alessio Siviglia, Gianna Nossa, Jessie Mosso, Eloise Mougel, Jamie Near, Thi Ngoc Anh Dinh, Omar Zenteno, Bernard Lanz, Thanh Phong Lê, Cristina Cudalbu.

Supervision: Cristina Cudalbu, Thanh Phong Lê.

## Ethics declarations

### Ethical approval

All animal experiments were conducted according to federal and local ethical guidelines, and the protocols were approved by the local Committee on Animal Experimentation for the Canton of Vaud, Switzerland (VD 3892).

