## Supplementary Information for "MRS4Brain: a processing toolbox for preclinical MR spectroscopy and spectroscopic imaging data"

### Installation guideline

The *MRS4Brain Toolbox* is compatible with MATLAB version 2023a and later across all major operating systems. Three MATLAB add-on toolboxes are required to execute the toolbox's processing pipelines: the Image Processing Toolbox (version 11.7 or newer), the Signal Processing Toolbox (version 9.2 or newer), the Parallel Computing Toolbox (version 7.8 or newer), and the Statistics and Machine Learning Toolbox (version 12.5 or newer).

In addition to MATLAB dependencies, four external software packages are necessary for full toolbox functionality: Advanced Normalization Tools (ANTs) for MRI co-registration, Docker to support the ANTs installation, LCModel for spectral fitting and quantification, and FID-A for specific preprocessing steps within the SVS pipeline. The FID-A functionalities required by the toolbox are already implemented internally, and therefore, no installation of FID-A is needed. Depending on the operating system, ANTsX/ANTs may require system-specific installation procedures for use in the MRI co-registration workflow. To streamline installation across platforms, including Windows and macOS, a Docker container (https://www.docker.com/) is employed. When the toolbox is used for the first time, if Docker is running, the toolbox automatically pulls the ANTs image into Docker. Users may alternatively download the image manually via Docker by searching for *antsx/ants:v2.5.0*. Download and installation may require several minutes to complete. Instructions for obtaining and installing LCModel are available on the LCModel website (https://s-provencher.com/lcmodel.shtml). Windows/macOS binaries are also provided in a GitHub repository with detailed installation guidance (https://github.com/schorschinho/LCModel).

At each new MATLAB session, directories for the toolbox folder and all its subfolders must be added to the MATLAB path. This can be accomplished by right-clicking the toolbox folder in MATLAB's Current Folder panel and selecting *Add to Path*, then *Selected Folders and Subfolders*, or by using the *genpath()* and *addpath()* commands in the Command Window. The toolbox is launched by entering *MRS4BRAIN_toolbox* in the MATLAB Command Window.

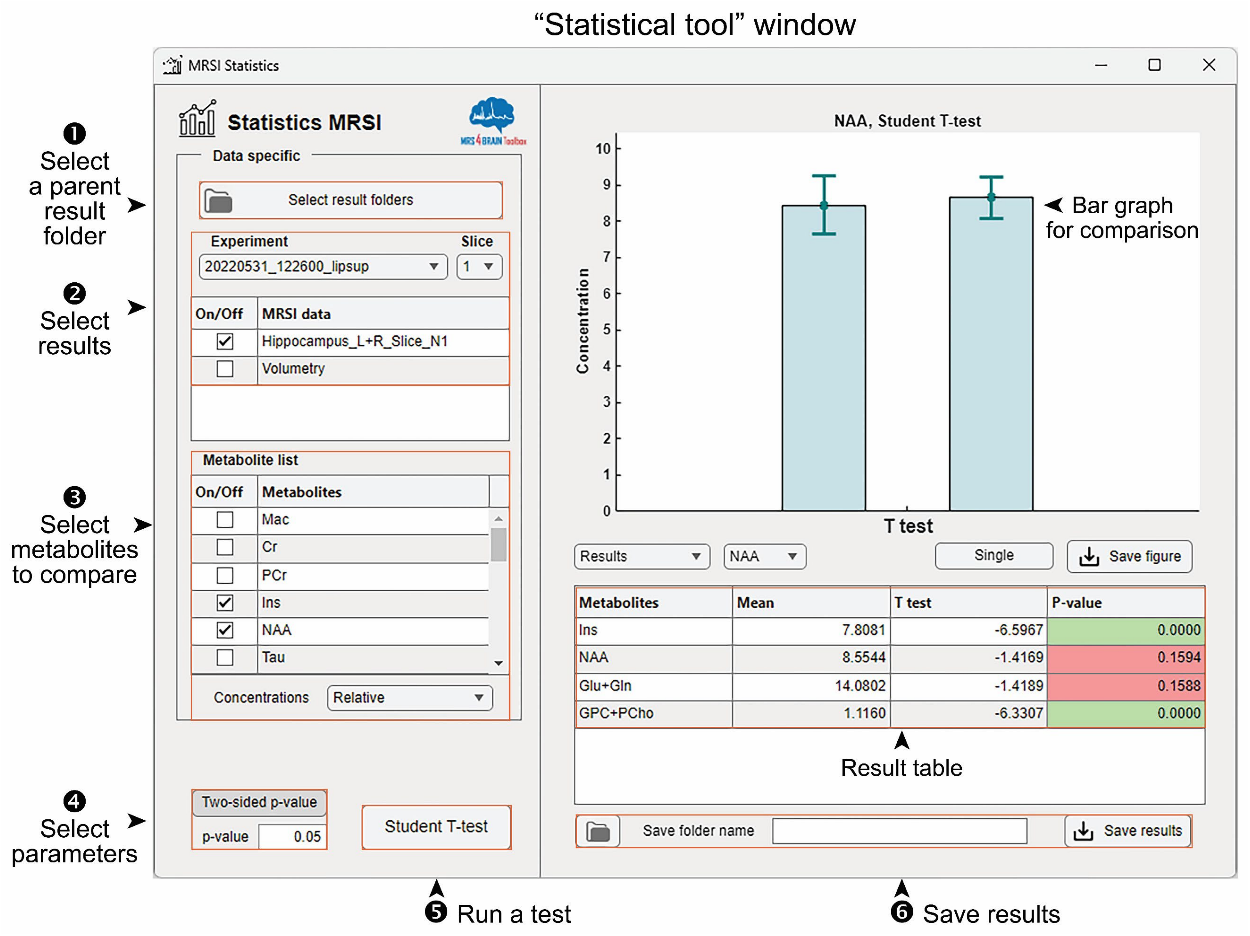


**Figure S1. Statistical tool for MRSI outputs.** Practical steps from importing results, selecting metabolites, the testing type, and its parameters, to performing a statistical analysis with graphs, *p*-values, and statistical values (e.g., *t*-value) provided for MRSI outputs.

**Supplementary Table 1**. Representative MRSI output: Hippocampus's concentration table of 9.4 T FID-MRSI data processed with no additional processing

| Metabolite | Mean relative concentration (a.u.) | Standard deviation (a.u.) | No. voxels |
|---|---|---|---|
| Mac | 0.0006 | 0.0001 | 46 |
| Cr | 3.7802 | 0.3605 | 37 |
| PCr | 4.3599 | 0.4819 | 42 |
| Ins | 8.0133 | 0.7241 | 45 |
| NAA | 10.2189 | 0.8034 | 46 |
| Tau | 5.8674 | 1.4941 | 46 |
| PCho | 1.7725 | 0.3297 | 8 |
| GPC | 1.5543 | 0.3268 | 3 |
| Glu | 9.7928 | 1.5043 | 46 |
| Gln | 2.9545 | 0.9671 | 39 |
| Ala | 2.5590 | 0.0000 | 1 |
| Asc | 4.2571 | 0.6209 | 38 |
| Asp | 4.9795 | 0.8754 | 36 |
| GABA | 2.6505 | 0.5938 | 25 |
| Glc | 5.7495 | 1.2408 | 28 |
| GSH | 1.9452 | 0.3363 | 36 |
| Lac | 2.6736 | 0.6741 | 35 |
| NAAG | 2.1719 | 1.0195 | 42 |
| PE | 0.0000 | 0.0000 | 0 |
| NAA+NAAG | 12.3467 | 1.5545 | 46 |
| Glu+Gln | 12.6337 | 2.2019 | 46 |
| GPC+PCho | 2.0451 | 0.3414 | 46 |
| Cr+PCr | 8.0000 | 0.0000 | 46 |

**Supplementary Table 2**. Representative SVS output: Concentration table of 9.4 T STEAM SVS data processed with spectral alignment and outlier removal

| Metabolite | Relative concentration (a.u.) | Absolute concentration (mmol/kg) | CRLB (%) |
|---|---|---|---|
| Mac | 0.000 | 0.000 | 2 |
| Cr | 4.025 | 4.362 | 4 |
| PCr | 3.975 | 4.308 | 4 |
| NAA | 8.937 | 9.686 | 2 |
| Tau | 6.697 | 7.258 | 3 |
| PCho | 0.635 | 0.688 | 24 |
| GPC | 0.405 | 0.439 | 36 |
| Glu | 9.011 | 9.767 | 3 |
| Gln | 3.204 | 3.473 | 6 |
| Ala | 0.623 | 0.675 | 20 |
| Asc | 2.622 | 2.842 | 9 |
| Asp | 1.642 | 1.780 | 19 |
| GABA | 1.079 | 1.170 | 14 |
| GSH | 0.846 | 0.917 | 13 |
| Lac | 2.035 | 2.206 | 7 |
| Ins | 6.839 | 7.412 | 3 |
| NAAG | 0.898 | 0.973 | 10 |
| PE | 1.623 | 1.759 | 13 |
| BHB | 0.120 | 0.130 | 92 |
| NAA+NAAG | 9.835 | 10.659 | 2 |
| Glu+Gln | 12.216 | 13.240 | 3 |
| GPC+PCho | 1.040 | 1.127 | 5 |
| Cr+PCr | 8.000 | 8.671 | 2 |

**Supplementary Table 3**. Representative DWS output: "Concentration" table of 9.4 T dSPECIAL DWS data (b-value 1) processed with ISIS handling, spectral alignment, and outlier removal

| Metabolite | Relative concentration (a.u.) | Absolute concentration (a.u.)* | CRLB (%) |
|---|---|---|---|
| Mac | 0.0015 | 0.0000 | 3 |
| Cr | 3.9010 | 0.0136 | 5 |
| PCr | 4.0990 | 0.0143 | 4 |
| Ins | 6.3830 | 0.0222 | 3 |
| NAA | 9.4710 | 0.0330 | 1 |
| Tau | 4.3300 | 0.0151 | 3 |
| PCho | 1.0310 | 0.0036 | 5 |
| GPC | 0.0000 | 0.0000 | 999 |
| Glu | 9.1380 | 0.0318 | 2 |
| Gln | 3.0320 | 0.0106 | 5 |
| Ala | 0.0850 | 0.0003 | 120 |
| Asc | 2.5410 | 0.0088 | 8 |
| Asp | 1.4730 | 0.0051 | 24 |
| GABA | 1.2110 | 0.0042 | 13 |
| Glc | 2.3680 | 0.0082 | 12 |
| GSH | 0.8150 | 0.0028 | 12 |
| Lac | 1.1020 | 0.0038 | 9 |
| NAAG | 0.3220 | 0.0011 | 29 |
| PE | 2.1230 | 0.0074 | 11 |
| NAA+NAAG | 9.7930 | 0.0341 | 1 |
| Glu+Gln | 12.1690 | 0.0423 | 2 |
| GPC+PCho | 1.0310 | 0.0036 | 5 |
| Cr+PCr | 8.0000 | 0.0278 | 1 |

* No reference water data is specified; therefore, absolute "concentration" values are in arbitrary units.
